# A diversity of diversities: do complex environmental effects underpin associations between below- and aboveground taxa?

**DOI:** 10.1101/2023.06.07.544003

**Authors:** Fiona M. Seaton, Paul B. L. George, Jamie Alison, Davey L. Jones, Simon Creer, Simon M. Smart, Bridget Emmett, David A. Robinson

**Author notes:** Corresponding author: Fiona M. Seaton, UK Centre for Ecology & Hydrology, Lancaster Environment Centre, Library Avenue, Bailrigg, Lancaster, LA1 4AP, UK.

## Abstract

1. To predict how biodiversity will respond to global change, it is crucial to understand the relative roles of abiotic drivers and biotic interactions in driving associations between the biodiversity of disparate taxa. It is particularly challenging to understand diversity-diversity links across domains and habitats, because data are rarely available for multiple above- and below-ground taxa across multiple sites.
2. Here we analyse data from a unique biodiversity dataset gathered across a variety of oceanic temperate terrestrial habitats in Wales, comprising 300 sites with co-located soil microbial, plant, bird, and pollinator surveys along with climate and soil physicochemical information. Soil groups are analysed using metabarcoding of the 16S, ITS1 and 18S DNA regions, allowing in depth characterisation of microbial and soil animal biodiversity.
3. We explore biodiversity relationships along three aspects of community composition: First, we assess correlation between the alpha diversity of different groups. Second, we assess whether biotic turnover between sites is correlated across different groups. Finally, we investigate the co-occurrence of individual taxa across sites. In each analysis, we assess the contribution of linear or non-linear environmental effects.
4. We find that a positive correlation between alpha diversity of plants, soil bacteria, soil fungi, soil heterotrophic protists, bees and butterflies is in fact driven by complex non-linear responses to abiotic drivers. In contrast, environmental variation did not account for positive associations between the diversity of plants and both birds and AM fungi, suggesting a role for biotic interactions.
5. Both the diversity and taxon-level associations between the differing soil groups remained even after accounting for non-linear environmental gradients. Aboveground, spatial factors played larger roles in driving biotic communities, while linear environmental gradients were sufficient to explain many group- and taxon-level relationships.
6. **Synthesis.** Our results show how non-linear responses to environmental gradients drive many of the relationships between plant biodiversity and the biodiversity of above- and belowground biological communities. Our work shows how different aspects of biodiversity might respond non-linearly to changing environments and identifies cases where management-induced changes in one community could either influence other taxa or lead to loss of apparent biological associations.

## Introduction

Ecosystem functionality and stability are underpinned by biodiversity (IPBES, 2019) and interactions between different components of the biosphere are essential to ecosystem maintenance. Correlation between the different biological communities of an ecosystem could be due to the direct influence of one taxonomic group on another or due to shared response to environmental conditions (Schulze et al., 2004; Wolters et al., 2006). Consequently, taxonomic or environmental mechanisms may lead to associated patterns of biodiversity but have differing implications for land use management. Indeed, opposing trajectories of above- and below-ground biodiversity as well as weakened biotic correlations have been observed in response to land use intensification (Gossner et al., 2016; Manning et al., 2015). Therefore, disentangling the relative influence of abiotic and biotic impacts upon the different components of the biosphere is key to understanding and predicting future responses of biological communities to environmental change.

There is a prevailing notion in ecology that diversity begets diversity. For example, a more diverse plant community can support a more diverse pollinator community through a proliferation of ecological niches (Schulze et al., 2004). There is still uncertainty over the strength of this synergistic effect, as relationships between the diversity of different taxa aboveground have been shown to be variable and weak (Wolters et al., 2006). Results investigating the relationship between above- and belowground biodiversity have also been variable, with either positive or no correlations commonly observed between plant diversity and soil microbial diversity (Hiiesalu et al., 2014; Leff et al., 2018; Prober et al., 2015). The different spatio-temporal scales of each study may explain some of these discrepancies, as the scale studied has been found to influence the relationships found between the diversity of different groups and their response to environmental factors (Chase et al., 2018; Wolters et al., 2006). Certain taxonomic groups appear to show more positive relationships with plant diversity. For example, fungal diversity, particularly mycorrhizal diversity, is often found to be positively related to plant diversity (Hiiesalu et al., 2014; Milcu et al., 2013; Nguyen et al., 2016; Peay et al., 2013; Ren et al., 2017; T. Yang et al., 2017). The type of plants that are considered in plant diversity inventories can also be important, for example the number of flowering plant species and their abundance has been shown to influence bee and butterfly abundance and richness (Kearns & Oliveras, 2009; Potts et al., 2009).

While relationships between the diversity of different taxonomic groups may be variable, there have been many results demonstrating that the species composition of plant assemblages may be related to the composition of other taxonomic groups even when diversity is not. Relationships between the community turnover of plants and the turnover of soil bacteria, fungi and protist communities have been found in both experimental and observational studies (Barberán et al., 2015; Cline et al., 2017; Delgado-Baquerizo et al., 2018; Leff et al., 2018; Prober et al., 2015). These correlations in composition and turnover in observational results can persist even after controlling for environmental drivers such as climate or soil physicochemical properties (Delgado-Baquerizo et al., 2018; Prober et al., 2015). Such changes in composition could be due to species-level associations between taxa, such as trophic interactions, symbioses or parasitic links.

Shared responses to environmental gradients could explain many of the measured associations between the biodiversity of different taxonomic groups. Some of the positive correlations that have been found between plant diversity and soil bacterial, fungal and protistan diversity are explained by shared response to environmental variables, e.g. soil pH and fertility (Goberna et al., 2016; Yashiro et al., 2018; Yuan et al., 2017). However, there are still many ecosystems, especially in extreme environments such as sub-polar regions and intensively managed agricultural systems where environmental factors may in fact cause divergent patterns and disconnectance of biodiversity (Cameron et al., 2019; Manning et al., 2015). Identifying the role of environmental factors in determining cross-taxa biodiversity associations can be challenging, with a vast array of potential environmental drivers that can each have direct, indirect, linear, non-linear, synergistic, and/or antagonistic effects on the species within an ecosystem. The scale of the environmental gradient also impacts the relative role of the environment in driving biodiversity change, as larger environmental gradients can have greater, or even non-linear, effects (Chase et al., 2018). Experimental manipulations, such as those established to evaluate the impact of plant diversity upon the biodiversity of the other groups, can yield useful information on the potential role of biodiversity interactions under relatively constant environmental conditions. Manipulative studies have found positive relations between plant diversity and the diversity of some other organisms, particularly for aboveground herbivores but less so for soil microbial groups (Cline et al., 2017; Dassen et al., 2017; Lind et al., 2015; Scherber et al., 2010; Weisser et al., 2017). However, by necessity empirical studies are limited in scope, covering a limited set of environmental conditions and plant species combinations. Furthermore, too few studies consider complex, non-linear environmental effects when assessing relationships between diversity or composition of disparate taxonomic groups.

Using a large multi-trophic field dataset spanning all terrestrial habitats in Wales, UK, here we attempt to unpick the relationships between a breadth of above and below-ground taxa. Specifically, we explore associations between plants, birds, pollinators and belowground soil bacteria, fungi (total fungi and arbuscular mycorrhizal fungi), heterotrophic protists and animals, both before and after accounting for linear or complex non-linear environmental effects. We assess the relationships between taxa across three axes of biodiversity, addressing aspects of alpha diversity, community turnover and taxon co-occurrence. We investigate (1) correlation between the alpha diversity of different groups using Bayesian hierarchical regression models; (2) whether turnover between sites is correlated across different groups using ecological distance analysis; and (3) the co-occurrence of individual taxa across sites, irrespective of higher groupings. In each analysis, we assess the contribution of linear or complex non-linear environmental effects. We hypothesise that (a) many inter- and intra-group relationships are readily explained by environmental variation, especially when complex, non-linear effects are considered; (b) those relationships remaining after accounting for environmental variation are more likely to be between groups and taxa that are known to have biotic interactions (e.g. nectar producing plants and pollinators, or herbaceous plants and soil fungi); and (c) that accounting for environmental effects would be more likely to cause associations between the alpha diversity of different groups to disappear compared to associations between community turnover or co-occurrence of individual taxa.

## Methods

### Field measurement programme

The data was collected as part of the Glastir Monitoring and Evaluation Programme (GMEP) field measurement program in Wales (Emmett et al., 2017; Wood et al., 2021). In total, 300 individual 1 km squares were randomly selected from within land classification strata in proportion to their extent across Wales in order to be representative of the range of habitat types across Wales or targeted to areas with high potential for agri-environment scheme uptake (Wood et al., 2021). Sampling occurred over a five-month period across each of the summers of 2013 to 2016; each square was only surveyed once over the four years with different squares being surveyed each year. Every square was subjected to a habitat survey, bird survey, two pollinator survey transects and multiple plant survey plots. A representative schematic of the survey layout within a 1km square is shown in Supplementary Fig. 1. For each square there were up to five 200 m^2^ square plant survey plots that also had soil samples taken. Soil samples were analysed for a variety of soil physicochemical properties, including pH in 1:2.5 CaCl_2_ suspension, total carbon, nitrogen and phosphorus, bulk density, water content, water repellency and electrical conductivity in 1:2.5 distilled water suspension. The data and full methods are available on the Environmental Information Data Centre (EIDC) (Robinson et al., 2019). Within the first two years of the survey soil samples were taken for microbial and eukaryotic community composition analysis from three of the 200 m^2^ plots, randomly selected per each 1 km square.

The habitats of each square were mapped and each plot was assigned to a habitat according to the UK Joint Nature Conservation Committee criteria (Jackson, 2000). The plot level measurements of microbial diversity were derived from 31% improved grassland; 23% neutral grassland; 12% acid grassland; 7% broadleaved woodland; 7% coniferous woodland; 7% bog; 4% arable; 3% dwarf shrub heath; 3% fen, marsh and swamp; and 2% bracken. Precipitation was the annual average rainfall and temperature was the annual average daily temperature for 1981-2010 calculated on a 1 km grid, this time period was chosen as it was the closest 30-year average provided by the Met Office that did not include any years post-survey. All climate data came from the Met Office © Crown copyright 2019 (Met Office et al., 2019).

### Biological data

#### Plant survey

Vegetation surveys were conducted for multiple plots per square, with the data being publicly available on the EIDC (Smart et al., 2020). Overall, there were ten types of sampling plots in total. Some plots were targeted to specific landscape features (e.g. hedges), meaning the total number and proportion of plot types varies across squares. The total number of vascular plant species recorded across the entire 1 km square was used as plant species richness for the square level analyses. Due to the differences in sampling effort across the squares, rarefaction curves were constructed to confirm that the number of plant species found had reached saturation. To assess whether different subsets of plants had greater associations with the diversity of other groups, within each plot the plant community was split by growth form, and for the aboveground communities the richness of plant species that were important to the diet of lowland birds, butterfly larvae and nectar provision was also calculated (Baude et al., 2016; Smart et al., 2000). Previous research has shown herbaceous plants to be more related to soil microbial groups than woody plants (Hu et al., 2018; Wang et al., 2016), however we did not expect all herbaceous groups (forbs, grasses, ferns, sedges, other monocots) to show the same relationships with soil microbial groups so we limit our herbaceous analysis to the forbs and grasses as they are the most diverse herbaceous groups within our dataset.

#### Bird and pollinator surveys

Birds were surveyed based on four morning visits to each square, equally spaced through mid-March to mid-July. Surveyors walked a route that passed within 50 m of all parts of each survey square, varying the start point for each survey in order to visit all parts of the square at least once before 08:00 h. All bird species seen and heard were recorded within each visit, and the total number of bird species recorded across all visits per square was used to calculate total bird species richness. Pollinator surveys were split into two independent parts: two 1 km transect routes separated by at least 500 m and where possible 250 m from the edge of the 1 km square; and a 20-minute timed search in a 150 m^2^ flower rich area within the 1 km square. The total number of butterfly species and bee groups recorded per square were used to calculate richness. Pollinator surveys were managed by Butterfly Conservation and bird surveys managed by the British Trust for Ornithology, and both datasets are publicly available on the EIDC (Botham et al., 2020; Siriwardena et al., 2020). Further details on survey methods are available within the data documentation upon the EIDC, and also within Wood et al. (2021).

#### Soil microbial diversity and community composition

Soil samples for microbial biodiversity analyses were taken using a gouge auger at 5 points around the physicochemical soil core location down to 15 cm, and then bulking together the samples. DNA was extracted from the combined, homogenised samples using mechanical lysis following homogenisation in triplicate from 0.25 g of soil per sample. The 16S (V4), ITS1 and 18S regions of the rRNA marker gene were targeted for amplicon sequencing to analyse the bacterial, fungal, and general eukaryotic diversity respectively. Diversity of arbuscular mycorrhizal fungi (AM fungi) was assessed separately using 18S data due to poor resolution with the ITS region (Berruti et al., 2017). For a full description of the methods used see George et al. (2019). Sequences with associated sample metadata have been uploaded to The European Nucleotide Archive with the following primary accession codes: PRJEB27883 (16S), PRJEB28028 (ITS1) and PRJEB28067 (18S).

Amplicon Sequence Variants (ASVs) were identified from the Illumina output using the DADA2 algorithm in R (Callahan et al., 2016). First, Illumina adapters were trimmed from sequences using Cutadapt (Martin, 2011). Then, the DADA2 package was used to filter all 16S, 18S and ITS1 reads to be truncated after the first instance of a quality score of 2, to remove all Ns and to have no more than 2 errors. Based on the examination of the change in quality scores with read length for a random subset of samples 16S reads were truncated to 200 bases, 18S reads truncated to 250 bases and ITS reads fewer than 100 bases in length removed. The DADA2 algorithm was then used to de-noise and merge the paired reads at standard settings. Sequence tables were constructed from the resultant ASVs and chimeric sequences were removed using default settings. Taxonomy was assigned using the DECIPHER R package (Wright, 2016), with both 16S and 18S reads being matched to SILVA r138 (Glöckner et al., 2017) and ITS being matched to UNITE v2019 (Nilsson et al., 2019). Non-bacterial ASVs were removed from the 16S data, and non-fungal ASVs from the ITS data. All ASVs appearing in the negative controls were removed from the analysis.

To account for the differences in read depth between samples, rarefaction was used as it has been shown to preserve the microbial relationships with biological origin (Weiss et al., 2017). The fungal data was rarefied to 10000 reads, bacteria to 20000 reads, and 18S eukaryotes to 20000 reads. Samples below this threshold were discarded, resulting in 437 fungal (ITS) measurements, 437 bacterial measurements and 438 eukaryote (18S) measurements. Eukaryote data was separated in to three sections based on higher order identification in the 18S dataset: AM fungi (phylum Glomeromycota), heterotrophic protists (Cercozoa, Ciliophora and Amoebozoa) and soil animals. Richness was calculated after rarefaction, which was repeated 100 times and the average result used in further analysis. For analysis of community composition the average of 20 rarefaction repeats was used.

### Statistics

#### Alpha diversity

All statistical analyses were performed within R version 4.3.0 (R Core Team, 2023). For analysis (1), assessing correlations between alpha diversity of different groups, we tested for effects of environmental variables using Bayesian multivariate hierarchical modelling within the brms package in R (Bürkner, 2017). This approach allowed us to include residual correlations between the alpha diversity of different groups, which would not have been possible if fitting multiple univariate models, without imposing specific directionalities of relationships within groups, as would be required using structural equation modelling. It also allowed us to include complex environmental effects, including non-linear effects, and a hierarchical effect of the 1km square for the plot-level models or of the vice county for the 1km square-level models. Group richness (for the plot-level model with soil groups and plant community) or group Shannon diversity (for the 1km square-level plant, bird and pollinator model) were modelled in a multivariate framework as either a function of a) solely the hierarchical spatial grouping, b) a linear function of abiotic variables, or c) as a non-linear function of abiotic variables.

For the plot-level model with linear abiotic drivers we included soil pH, total soil carbon (log transformed), water content, total phosphorus and soil water repellency (log transformed) as predictors, with a unit normal prior on all effects. These environmental predictors were chosen due to previous analysis of this dataset showing that the effect of climate upon the soil microbial communities was mediated by changes in soil physico-chemical properties (Seaton et al., 2019, 2020). Bulk density and soil nitrogen were not included in the model due to being highly correlated with soil carbon (correlation coefficients of -0.92 and 0.92 respectively), without these variables the highest absolute correlation was 0.6 which we judged not to be an issue for model fitting. Habitat type is not used as a predictor in any of these models, as our habitat data was determined by examining the plant communities which leads to a circularity problem where we would be predicting the plant community using data derived from the plant community. When including non-linear environmental effects, we limited the model to only a sigmoidal relationship between microbial richness and pH due to the pre-existing literature on the non-linear effects of soil pH on soil biochemical properties (Aciego Pietri & Brookes, 2008; Bickel et al., 2019; Lauber et al., 2009). A sigmoidal model was chosen rather than a bell curve as there were too few soils in the pH > 7 range, where bacterial richness has been shown to decline, to estimate any such decline. No other environmental variables were included as predictors, as we wanted to test the hypothesis that a single, highly influential, variable that is appropriately fully accounted for through modelling a non-linear effect could be as important as including multiple environmental variables but only modelling them as linear effects. This sigmoidal model took the form shown in equation 1, where parameters α (upper asymptote), β (growth rate), γ (x value at maximum growth), and δ (intercept) were fit using weakly informative priors.

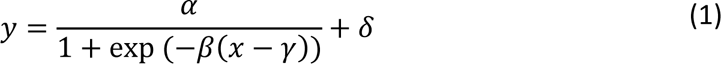

For the square-level model with linear abiotic drivers we included annual temperature and precipitation as predictors, again with a unit normal prior on all effects. We again did not use habitat type as a predictor to avoid the circularity issue described above. To investigate a role for complex environmental effects, the plant, bird and pollinator model with linear environmental effects was compared to a model where the exponentiated Shannon diversity values were modelled as a function of interacting temperature and precipitation.

The square identity was included as a group-level effect on the intercept (δ) in order to account for the spatial element of the data. To evaluate whether associations between the alpha diversity of the different groups remained after accounting for environment the residual correlations between groups were extracted from the models. All response variables were modelled as normal, to allow for both the rarefaction process for microbial groups detailed above resulting in non-integer responses and the inclusion of residual correlations within the model structure. Satisfactory recovery of the data was confirmed using graphical posterior predictive checks. No correlations between the group-level effects were included within the model.

#### Community turnover

For analysis (2), assessing the correlations in turnover between sites across different groups, we carried out an ordination analysis using non-metric dimensional scaling (NMDS). This allowed us to move beyond simple summary metrics such as alpha diversity and instead consider associations in broad-scale compositional changes across the different groups. Distance calculation and NMDS were performed using the vegan package (Oksanen et al., 2020). Binary Jaccard distance was used for plant, bird and butterfly species composition and Bray-Curtis distance used for bee and hoverfly groups due to their lower richness. All soil microbial groups were compared based upon Bray-Curtis distance. Direct correlation coefficients between the turnover of the different groups were calculated using Procrustes analysis upon the PCoA scores for each group. Forbs, grasses and AM fungi did not appear in every plot, so those correlations are based upon a subset of the data, woody species appeared in too few plots (36%) to be included. To unpick the relative contribution of shared environmental drivers, interaction with the plant community and spatial autocorrelation that could be driving changes in the main axes of variation in the communities the variance in the ordination scores of each group was partitioned using the varpart function within the vegan package. The response variables were the NMDS scores of each group based on Jaccard distance for aboveground groups and Bray-Curtis distance for belowground groups. The NMDS for the plot-level analysis were mostly performed using 3 dimensions, and had stress values of 0.067 for bacteria, 0.117 for fungi, 0.095 for heterotrophic protists, 0.192 for soil animals, and 0.076 for plants (over 4 dimensions). No stable NMDS solution was found for AM fungi even after varying the number of dimensions, ecological distance used, and convergence criteria so they were omitted from this analysis. The NMDS for the square level analysis were mostly performed using 4 dimensions and had stress values of 0.098 for plants, 0.089 for birds, 0.139 for bees and hoverflies (3 dimensions), and 0.139 for butterflies. The predictors within the variance partitioning were climate (represented by temperature and precipitation), spatial distance (represented by principal coordinates of neighbour matrices), plant community composition (represented by the NMDS scores). For the plot level data only the first four dimensions of a PCA upon soil physicochemical properties (pH, carbon, nitrogen, total phosphorus, bulk density, electrical conductivity, water content, water repellency) were also used as predictors. These dimensions explained a total of 88% of the variance in the measured soil physicochemical properties.

#### Taxon-level relationships

For analysis (3), assessing co-occurrence of taxa irrespective of higher groupings, relationships were evaluated using joint species distribution models to identify whether co-occurrences could be explained by shared responses to environmental drivers and spatial autocorrelation. This approach allowed us to move further from the broad-level compositional associations in analysis (2) by modelling each individual species separately, allowing us to evaluate both intra- and inter-group species-level associations within the same framework. Joint species distribution models were fitted using the sjSDM package in R (Pichler & Hartig, 2021). Only common taxa were included within the analysis to reduce the number of false positive associations (Weiss et al., 2016), which we define as being present in 25% of sites for bacteria, heterotrophic protists, fungi and soil animals and 10% for plants, birds and pollinators. In total there were 2309 bacterial taxa, 236 heterotrophic protists, 190 fungal taxa, 22 animals and 28 plants included in the plot-level belowground analysis while the square-level aboveground analysis included 348 plant species, 101 bird species, 22 butterfly species, 4 bee groups and 4 hoverfly groups. All models had the spatial relationships between sites modelled as a linear function of the spatial eigenvectors, all taxa converted to presence/absence and fit with a binomial model with probit link, and all other parameters set to their defaults. The responses to environmental gradients were either excluded (intercept-only model) or modelled as either linear responses or deep neural networks. Within the plot-level model (i.e. including microbial groups and plant measurements from individual plots) the environmental predictors were soil pH, organic matter, carbon, nitrogen, available phosphorus (Olsen-P), conductivity, water repellency, moisture content and bulk density. Within the aboveground model (i.e. including plants at the square level, birds and pollinating insects) the environmental predictors were average annual precipitation (1981-2010), daily temperature (1981-2010) and elevation.

## Results

### Alpha diversity

We found that bacterial, fungal, heterotrophic protistan, bird, bee and butterfly richness were all positively correlated with plant diversity. Within the soil groups, AM fungi showed the strongest correlation with plants, whereas bacteria, general fungi and heterotrophic protists all had lower correlations, and no correlation with animal richness (Table 1, upper triangle). All soil groups were positively correlated with each other; however animal richness was only weakly associated with the other groups. Soil microbial richness was more strongly positively correlated to the richness of forb plant species as opposed to overall plant or grass species richness. Microbial richness was also negatively correlated to woody plant richness, particularly in AM fungi (ρ = -0.65) and heterotrophic protists (ρ = -0.51). Plant diversity in the 1 km squares positively correlated most strongly with bird species richness (ρ = 0.50), followed by butterfly species richness (ρ = 0.43), then bee group and hoverfly group (ρ ∼ 0.35) richness. Limiting the plant community to those species known to be important to the different aboveground animal groups through food provision resulted in an increase in correlation coefficients of 0.08 for birds but not for the other groups. The different pollinator groups were positively associated with each other, with butterflies and bees associated more strongly with each other than hoverflies with the other two groups (ρ = 0.56 compared to ρ 0.42-0.44). Bird species richness was positively correlated with butterfly and bee richness (ρ ∼ 0.5). Average soil microbial diversity per 1 km square was also positively correlated with bird, butterfly and hoverfly richness, with correlations ranging from ∼0.35 to ∼0.55.

**Table 1:** The correlations between the different microbial groups and the plant community (raw data, not model results), the upper triangle shows the Spearman rank correlation coefficient for alpha diversity (richness) and the lower triangle shows the Procrustes correlation coefficient for compositional turnover (Bray-Curtis distance).

|  |  |  |  |  |  |  |  |
| --- | --- | --- | --- | --- | --- | --- | --- |
| **Plant** | 0.22 | 0.21 | 0.19 | -0.02 | 0.40 | 0.81 | 0.69 |
| 0.54 | **Fungi** | 0.51 | 0.78 | 0.20 | 0.53 | 0.42 | 0.25 |
| 0.50 | 0.71 | **Bacteria** | 0.56 | 0.07 | 0.32 | 0.39 | 0.16 |
| 0.56 | 0.72 | 0.78 | **Hetero. protists** | 0.13 | 0.54 | 0.45 | 0.26 |
| 0.41 | 0.45 | 0.50 | 0.52 | **Soil animals** | -0.05 | -0.16 | -0.05 |
| 0.43 | 0.54 | 0.52 | 0.58 | 0.35 | **AM Fungi** | 0.57 | 0.54 |
| 0.55 | 0.47 | 0.46 | 0.51 | 0.32 | 0.40 | **Forbs** | 0.56 |
| 0.92 | 0.47 | 0.49 | 0.52 | 0.36 | 0.42 | 0.46 | **Grasses** |

#### The effects of environmental factors upon alpha diversity relationships

Accounting for environmental gradients reduced the magnitude of the correlations between the diversity of the different taxonomic groups both above and belowground (Fig. 1). Aboveground, only the correlation between plant diversity and bird diversity and the correlation between bee diversity and butterfly diversity remained after accounting for the temperature and rainfall gradients (Fig. 1a-c). It might be expected that the diversity of plant groups that are important to the diets of the different aboveground animal groups would show stronger relationships with these groups, and this was tested by comparing direct models of plant richness as a predictor of bird, butterfly and bee richness versus models with the richness of plants important in the diet of lowland birds, butterfly larval food plants and nectar producing plants respectively. However, we found no improvement in model predictive performance compared to models with overall plant richness.

**Figure 1:**
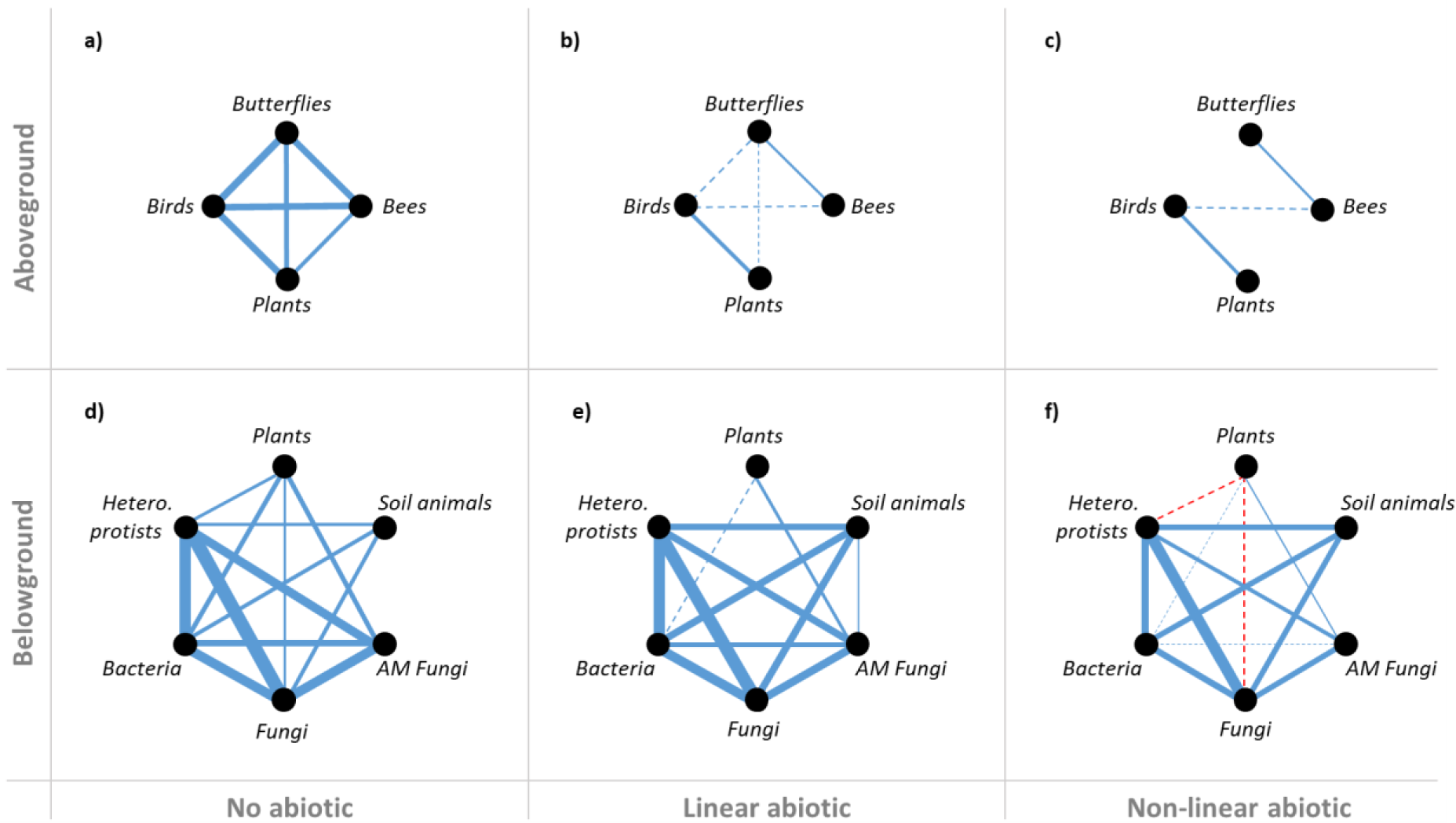
The residual correlations between the diversity of the different groups if no abiotic drivers are taken into account (a, d), if abiotic drivers are included as linear effects only (b, e), and if non-linear effects are included (c, f) are included. Blue lines indicate positive residual correlations, red lines negative. The width of each line is proportional to the estimated residual correlation, lines are solid if the 95% interval did not include zero or dashed if the 95% interval included zero but the 60% interval did not. Residual correlations where the 60% interval included zero are not shown in the figure but are included within the model. For the full model summary see Supplementary tables 1-6.

Belowground, the richness of the soil groups did maintain correlations with each other even after accounting for changes in soil properties, and in the case of soil animal richness the residual correlation with the other groups actually increased once a variety of soil properties including soil carbon were accounted for (Fig. 1d-e, Supplementary tables 1-2). For all other soil groups, the model with a sigmoidal pH gradient accounted for more residual correlation (Fig. 1f) than the linear models of multiple soil properties (Fig. 1e). All soil microbial groups, in addition to plant richness, showed a good fit of the sigmoidal pH model to the data while soil animal richness showed a linear decrease with increasing pH (Fig. 2). Accounting for soil properties removed the residual correlation between plant richness and soil richness for all soil groups other than AM fungi, which retained a positive relationship with plant richness.

**Figure 2:**
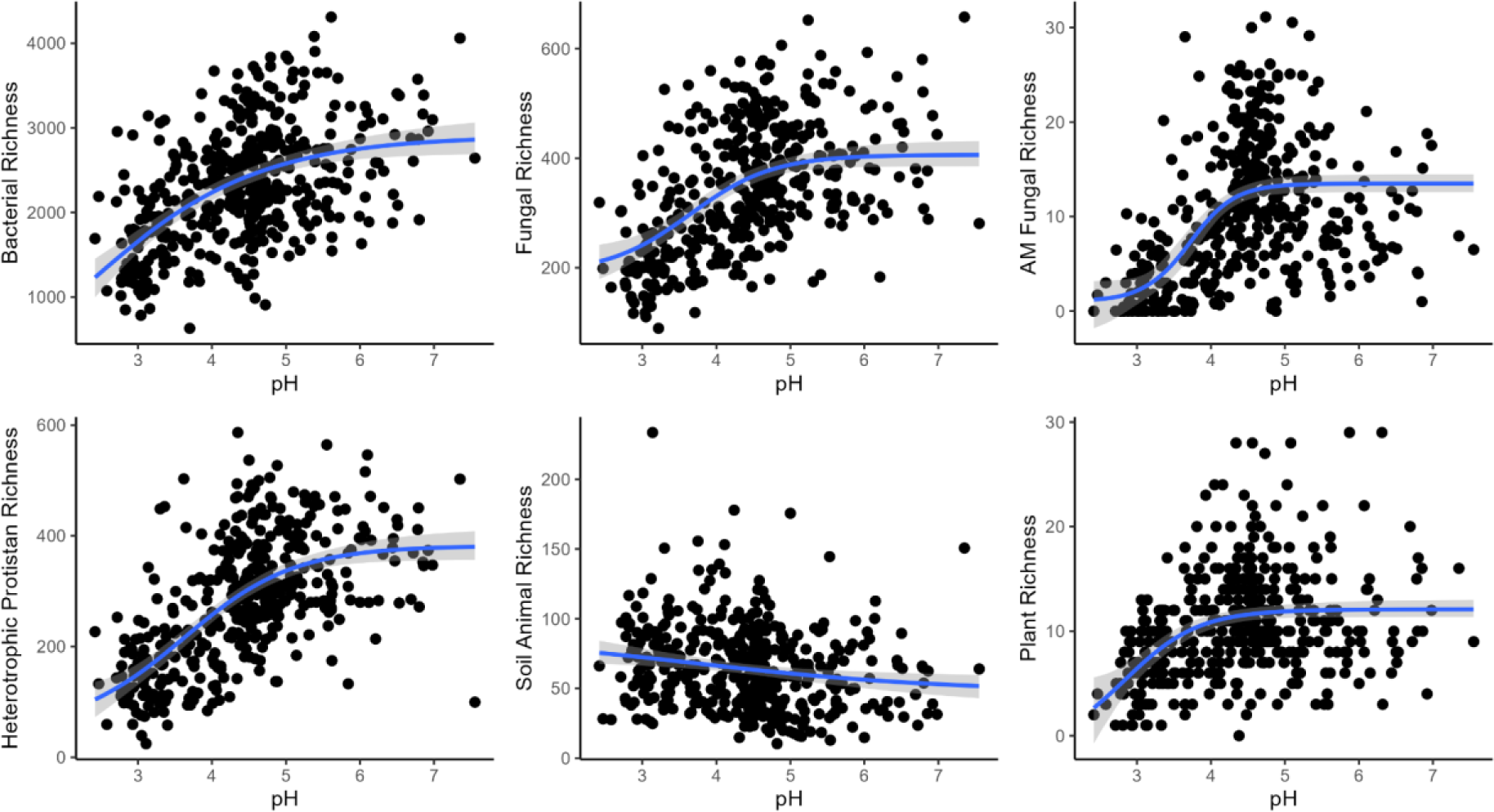
Predicted effect of soil pH on soil and plant diversity in the multivariate model with a sigmoidal pH effect. Plots show the actual values as black dots and the conditional pH effect as a blue line with the 95% interval of the effect as a shaded grey ribbon.

### Community turnover

Plant community turnover was positively associated with fungal, bacterial, heterotrophic protistan, bird and butterfly community turnover. Bacteria, heterotrophic protists and general fungi showed a slightly stronger correlation with plant community turnover than AM fungi did (Table 1, lower triangle); all were significant at p = 0.001. Turnover in heterotrophic protistan communities strongly tracked turnover in bacterial communities, with a Procrustes correlation statistic of 0.78. Fungal community turnover also correlated with bacterial turnover, with a Procrustes correlation statistic of 0.71. Bird and butterfly community turnover were more strongly related to plant community turnover than the link between bee, hoverfly and plant turnover. This was reflected by both the spearman rank correlations and the Procrustes correlation statistic (0.73, 0.59, 0.23 and 0.32 for bird, butterflies, bees and hoverflies respectively). However, all the Procrustes analyses were found to be significant at p = 0.001.

#### The effects of environmental factors upon community turnover relationships

Plant community turnover explained unique variance in the bacterial, fungal and heterotrophic protist communities even after accounting for spatial and environmental factors (Fig. 3a). Plant community turnover at the plot scale was partially explained by soil, climate and spatial factors but had considerable unexplained variance (71%). Climate was initially included but did not explain any unique variance in turnover of the soil microbial communities and so was omitted from this analysis. The pattern of variance explained was similar across the soil groups, with the soil physicochemical properties and plant community jointly explaining more variation than the distance between samples. However, there was a large proportion of variation in fungi, bacteria, animals and heterotrophic protists that was explained by soil, plant and spatial factors together.

**Figure 3:**
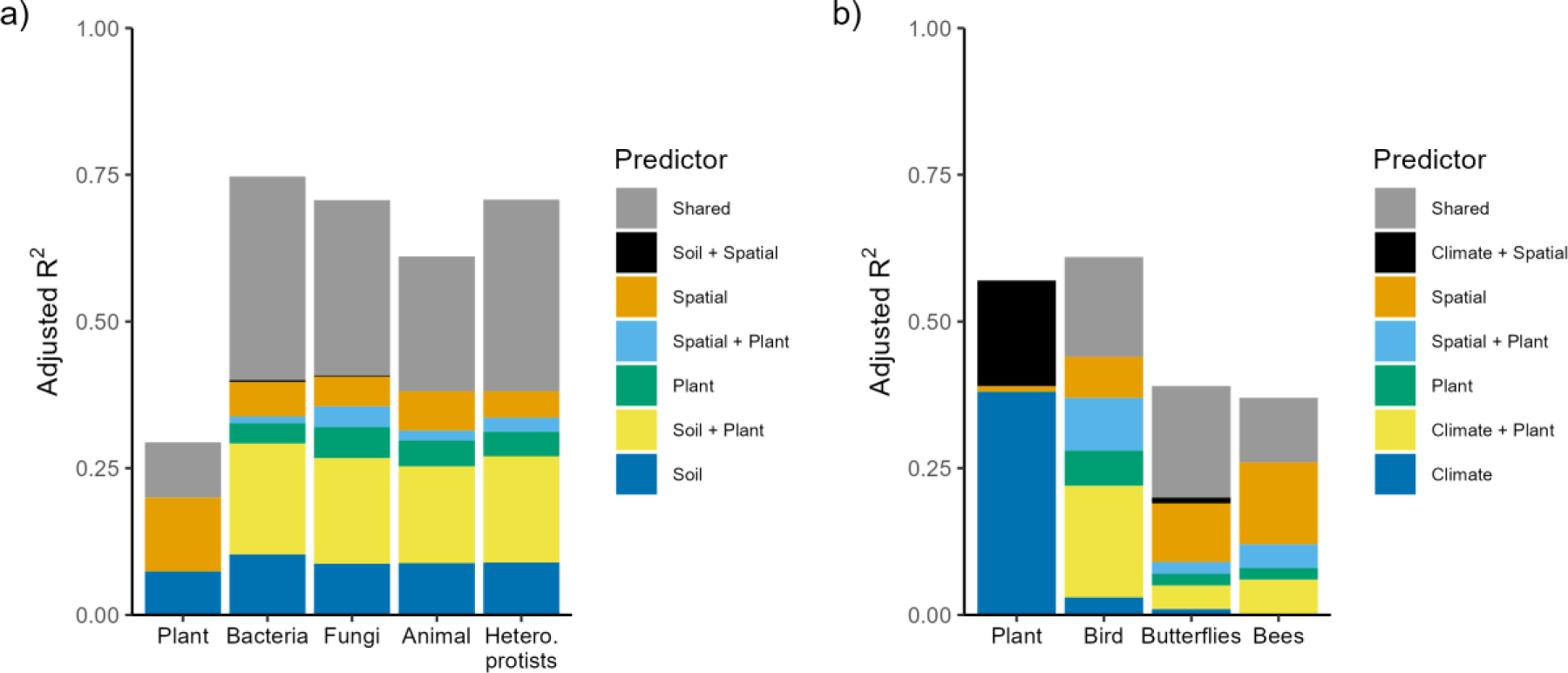
The properties that explain the variation in community turnover at the plot level (a) and the square level (b). The height of each coloured bar section represents the proportion of variation explained by the category, with the “Shared” category representing the variation explained by all factors together in addition to the variation explained by each single or two-part combination of factors.

Plant community turnover explained some of the variance in bird, butterfly and bee and hoverfly community turnover (Fig. 3b). Plant community turnover at the 1 km square level was largely explained by variation in precipitation and temperature (38%), with some effect of the distance between squares interacting with climate (19%). Turnover in the bird community was explained comparatively well by the plant community and the climate (28% without any spatial element), whilst also incorporating a spatial interaction between the two. The pollinator communities were explained relatively poorly; plant community turnover, climate and spatial factors did appear to be important together but overall ∼60% of variation was unexplained. Precipitation and temperature were less important to bees and hoverflies than they are to the other groups, with precipitation and temperature in general being the least important factor for the aboveground animal groups but much more important for plants. Spatial factors appeared to be relatively more important for the animal groups than they were for the plants.

### Taxon-level relationships

Residual co-occurrence relationships between common below- and aboveground taxa were identified by joint species distribution modelling both with and without accounting for shared responses to environmental variables. In all cases the average magnitude of taxon correlations was decreased once environmental gradients were accounted for, with non-linear environmental gradients removing more apparent species correlations than linear environmental gradients (Fig. 4). The effect of incorporating environmental conditions was greater upon apparent negative associations between taxa, with proportionally more negative associations lost after changes in environmental conditions were accounted for. Within the soil groups more positive associations were retained after accounting for environmental conditions, with the mode of the correlation distribution remaining above zero compared to it being centred on zero in all plant-related and aboveground correlation distributions. Soil groups showed on average differing levels of association with differing plant growth forms, with forbs showing more positive associations with soil microbial groups than grasses. However, accounting for non-linear environmental conditions removed the strength of the forb relationship in average correlation magnitude between plant growth forms (Supplementary Fig. 2a). Only one woody species (*Vaccinium myrtillus*) appeared in enough plots to be included in this analysis, and so those results are unlikely to be representative of all woody species. There was very little difference between the correlation strength of plants that are, or are not, food sources for birds and pollinators across the different models, with only a slight tendency to have a higher average magnitude of correlation for food source plants in the no environment and non-linear models but not in the non-linear model (Supplementary Fig. 2b).

**Figure 4.**
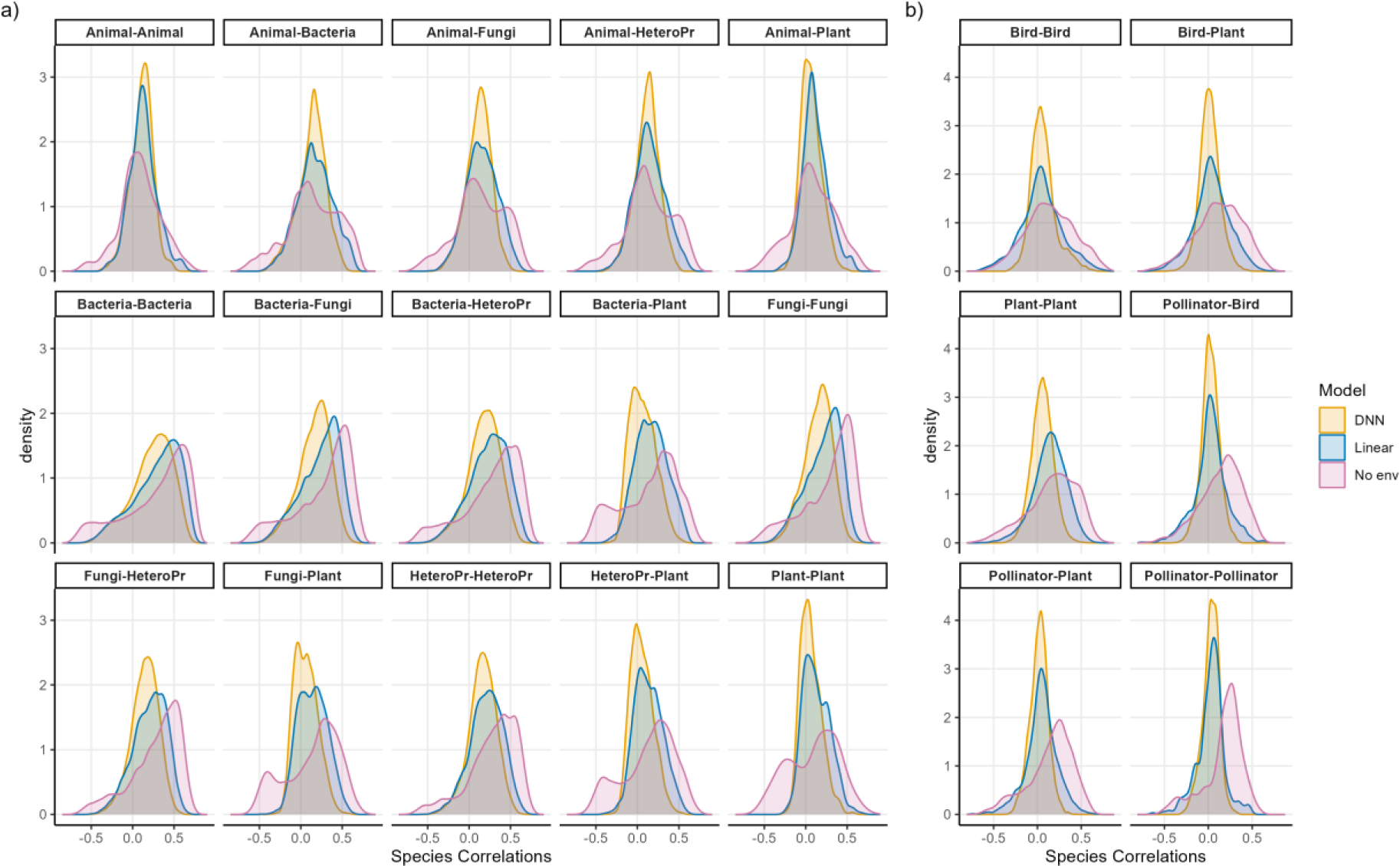
The density distributions of taxon correlations from the belowground/plot-level (a) and aboveground/square level (b) joint species distribution models that took no account of environment (No env), linear functions of environment only (Linear), or that used deep neural nets to allow for non-linear effects of environment (DNN). Butterflies, bees and hoverflies are combined into a pollinator category for graphical simplicity, as are general fungi and AM fungi.

Despite the overall trend towards lower average magnitude of species correlations after accounting for environmental conditions there was a small proportion of species correlations that actually increased in magnitude after accounting for environmental gradients. Around 7% of correlations increased above- and belowground by over 0.05 units in the non-linear environmental model compared to the model with only spatial and biotic components. The interactions between soil animal and other soil taxa showed proportionally more correlations that increased, with 17% of correlations increasing by >0.05, while the interactions between the plant and soil taxa showed proportionally more decreases in magnitude. The relative importance of biotic factors, environmental factors, and spatial factors varied considerably by taxa, although on average biotic factors appeared more important belowground than aboveground while spatial factors were more important aboveground than belowground (Supplementary Figs 3 and 4).

## Discussion

Consistent with our hypothesis we find that many alpha diversity relationships dissolve after accounting for abiotic effects, with non-linear environmental gradients clearly affecting belowground alpha diversity relationships. Our results indicate that the positive correlation of plant and soil microbial diversity is driven by changes in the soil environment and are in agreement with previous work showing the importance of soil pH and fertility in jointly influencing plant and microbial diversity (Goberna et al., 2016; Tedersoo et al., 2016; Yashiro et al., 2018; Yuan et al., 2017). Importantly, accounting for the influence of soil pH upon microbial diversity in a non-linear way led to the loss of a plant ∼ microbe diversity relationship, with the exception of AM fungi. AM fungi are more dependent upon plant inputs and are capable of growing through large volumes of soil, meaning that they would be expected to show stronger relations with the plant community (Hiiesalu et al., 2014; Nguyen et al., 2016). It is possible, however, that our models are attributing associations between groups to environmental factors when biotic interactions are occurring, particularly when biotic associations are in part driven by the environment. Therefore, our results provide a starting point for further investigation and identify those associations that seem most likely to be unrelated to shared environmental responses rather than providing a definitive answer as to the presence or absence of biotic associations. Our results also indicate a strong linkage between bacterial, fungal and heterotrophic protistan communities that could potentially represent a greater role of trophic dynamics relative to edaphic properties and plant inputs in structuring those communities.

The diversity of the pollinator groups showed no residual correlation with plant communities after accounting for environmental gradients, and in contrast to our hypothesis we found no evidence of stronger links between foodplants and the aboveground animals that feed on them compared to overall plant diversity. The lack of relationship between pollinator and plant diversity was unexpected due to the role of plants in providing food resources and niche space, and we found no evidence of stronger associations in what should be trophically interacting species at either the alpha diversity or the taxon-level. It could be that our environmental variables are taking up some of the variation explained by true biotic associations, however in these models we only included climatic data as environmental predictors which seems unlikely to be able to account for all plant-pollinator associations. However, previous research has shown variable relationships between plant and pollinator diversity, potentially related to vegetation structure and the greater importance of floral abundance and seasonality compared to plant or food-plant diversity (Alison et al., 2022; Berg et al., 2019; Lowe et al., 2022; Potts et al., 2009). We also only considered overall plant diversity and the diversity of food plants; the structural heterogeneity of the plant community could instead be the important driver of any plant-pollinator diversity relationships. In contrast, increasing bird diversity in conjunction with plant diversity was only partially confounded by the climate gradient, indicating that plant diversity could potentially have a direct impact upon bird diversity. This could be due to the provision of a greater variety of food sources, through increased landscape heterogeneity or potentially through the influence of an unidentified confounding variable such as anthropogenic activity (Le Provost et al., 2021; Stein & Kreft, 2014).

We found that the composition of above- and belowground communities was influenced by plant community composition even after accounting for environmental gradients, in contrast to the alpha diversity results. The proportion of variance in community turnover explained by plants was much higher in the bacterial, fungal and heterotrophic protist communities than in the bird, butterfly, bee and hoverfly communities. The finding that plant community turnover explains bacterial, fungal and heterotrophic protistan community turnover is consistent with previous findings at the field (Leff et al., 2018), regional (Barberán et al., 2015; Chen et al., 2017; T. Yang et al., 2017), and global scales (Delgado-Baquerizo et al., 2018; Prober et al., 2015). The higher proportion of explained variance in bird communities compared to butterfly, bee and hoverfly communities was unexpected as pollinators have in general been found to be closely related to plant composition (Hofmann et al., 2017; Kearns & Oliveras, 2009; Weisser et al., 2017). Overall, the majority of the variation in butterfly, bee and hoverfly community turnover was unexplained by climate, plant communities or spatial factors. This may be due to anthropogenic influences or other factors not incorporated into our model such as agri-environment scheme uptake, pollution, or habitat heterogeneity being important in driving pollinator community assembly.

The closer linkages of the soil groups to each other compared to the plant linkages above- and belowground were apparent in both the overall diversity and species-level results. Some of these results, particularly the alpha diversity results, may be due to the soil groups all being surveyed using a metabarcoding approach as opposed to the aboveground groups having their own dedicated survey methods. Processes such as DNA extraction efficiency could vary systematically across the plots, which we might expect to be related to changes in soil physico-chemical properties (e.g. acidity) and so accounting for environmental variables could help remove some residual metabarcoding-related associations. In particular, the soil animal community showed stronger linkages with other soil groups after accounting for environmental gradients. Soil animal groups are strongly influenced by soil carbon and other environmental factors (Caruso et al., 2019; George et al., 2017; Lavelle et al., 2022), and it appears that once these are accounted for the secondary role of biotic associations in driving community assembly becomes clearer. Some of these identified linkages between the taxa within and between groups may represent shared response to environmental or geographic drivers not included within our analysis, while some may instead represent true interactions across domains (Blanchet et al., 2020; Carr et al., 2019; Seaton et al., 2020).

The different taxonomic groups within our analysis operate upon different spatial and temporal scales, and this appears to have affected the relative influence of spatial factors upon biological communities that we have found (Chase et al., 2018). In our results we found that spatial distance appeared to have a greater influence upon the species composition of the aboveground communities measured at the 1km square level. Previous work has shown that the scale at which biodiversity is measured, impacts the strength of the correlation found, with 10 km^2^ areas showing the strongest relationship for aboveground diversity (Wolters et al., 2006). However, the life history traits of the community in question should be expected to impact the scale of interest, and within our dataset we have organisms that are whose mobility is limited to very short distances such as soil bacteria (P. Yang & van Elsas, 2018) ranging up to bird species that can travel vast distances. Therefore, the spatial factors we measured may not have been at the appropriate scale to influence the soil microbial communities at the plot level and the stronger relationships between the soil microbial groups may be related to the plot scale being more equivalent to the larger (>10 km^2^) scales that show greater associations aboveground. It is also worth considering the role of the spatio-temporal scale of survey in determining the range of the environmental drivers of biodiversity (Chase et al., 2018). Larger areas contain a wider range of environmental conditions where different parts of the range could have contrasting effects on the biosphere. Our survey covers a range of environmental conditions that is representative of the Welsh landscape, however it only contains a limited sampling of feasible potential environmental conditions such as intense agricultural landscapes more common in other areas of Europe where we might expect different influences upon soil and aboveground communities (Billeter et al., 2008; Gossner et al., 2016).

We found that accounting for non-linear environmental effects had a greater influence on inferred biotic correlations at both the overall diversity and species level compared to only incorporating linear environmental effects. This emphasises the importance of considering nonlinear effects to prevent underestimation of the impacts of environmental gradients as well as increase the applicability of the results outside the environmental range of each study, particularly within soil communities (Austin, 2007). In contrast, including only linear effects of the environment proved sufficient to remove most of the aboveground diversity and positive species associations. However, it is possible that other environmental drivers or landscapes may show a greater role of non-linear environmental effects in driving plant, pollinator, and bird community dynamics. It is also possible that changes in land use intensity or other factors could change the very nature of the relationships between the diversity and composition of differing components of the ecosystem (Felipe-Lucia et al., 2020; Le Provost et al., 2021).

Within this work we take a step towards disentangling the relative cross domain biotic and environmental associations governing community assembly across the terrestrial-soil habitat boundary. We have identified associations between groups that are less likely to be driven by environmental changes and thus more likely to be jointly disrupted by management changes, such as the relationships between plants and AM fungi where the sowing of more diverse plant communities could result in direct changes in AM fungi, or the relationships between soil animals and soil microbial groups where the loss of soil animals through pesticide application could disrupt the whole soil food web. We have also identified associations that do appear to be driven by environmental changes, and where direct changes to one group may result in disconnectance from the other groups, such as potentially between plants and pollinators which suggest that the pollinator community may not always be able to adapt to management-induced changes in the plant community due to the influence of climate. Importantly, our results provide insight into the importance of non-linear environmental effects upon ecological communities across a wide variety of entwined habitats and environmental conditions and provide potential future avenues of research in evaluating whole-ecosystem state and change, in relation to complex environmental drivers.

## Acknowledgements

We thank all the GMEP team who contributed to collecting all the data. The research was funded by the Welsh Government through the Glastir Monitoring and Evaluation Program (GMEP) with underpinning funding from the Natural Environment Research Council award number NE/R016429/1 as part of the UKScaPE programme delivering National Capability. Contract reference: C147/2010/11. NERC/Centre for Ecology & Hydrology (CEH Projects: NEC04780/NEC05371/NEC05782). F.M.S. and P.B.L.G were supported by the award of studentship grants by the NERC and BBSRC funded STARS (Soils Training And Research Studentships) Centre for Doctoral Training and Research Programme; a consortium consisting of Bangor University, British Geological Survey, Centre for Ecology and Hydrology, Cranfield University, James Hutton Institute, Lancaster University, Rothamsted Research and the University of Nottingham. Grant Ref: NE/M009106/1.

## Author contributions

Seaton, George, Creer, Jones, and Robinson conceived the ideas and Seaton designed the statistical methodology with input from Alison, George and Smart; Emmett and Robinson led the collection of the data; Seaton analysed the data and led the writing of the manuscript. All authors contributed critically to the drafts and gave final approval for publication.

## Data availability

The data that support the findings of this study are openly available, with the soil physicochemical data at DOI: 10.5285/0fa51dc6-1537-4ad6-9d06-e476c137ed09, the vegetation data at DOI: 10.5285/71d3619c-4439-4c9e-84dc-3ca873d7f5cc, the pollinator data at DOI: 10.5285/3c8f4e46-bf6c-4ea1-9340-571fede26ee8, the bird data at DOI: 10.5285/31da0a94-62be-47b3-b76e-4bdef3037360 and the soil microbial sequences data in the European Nucleotide Archive with primary accession codes: PRJEB27883 (16S), PRJEB28028 (ITS1) and PRJEB28067 (18S).

## Conflict of Interest

The authors have no conflicts of interest to declare.

## Supplementary Information

**Supplementary figure 1.**
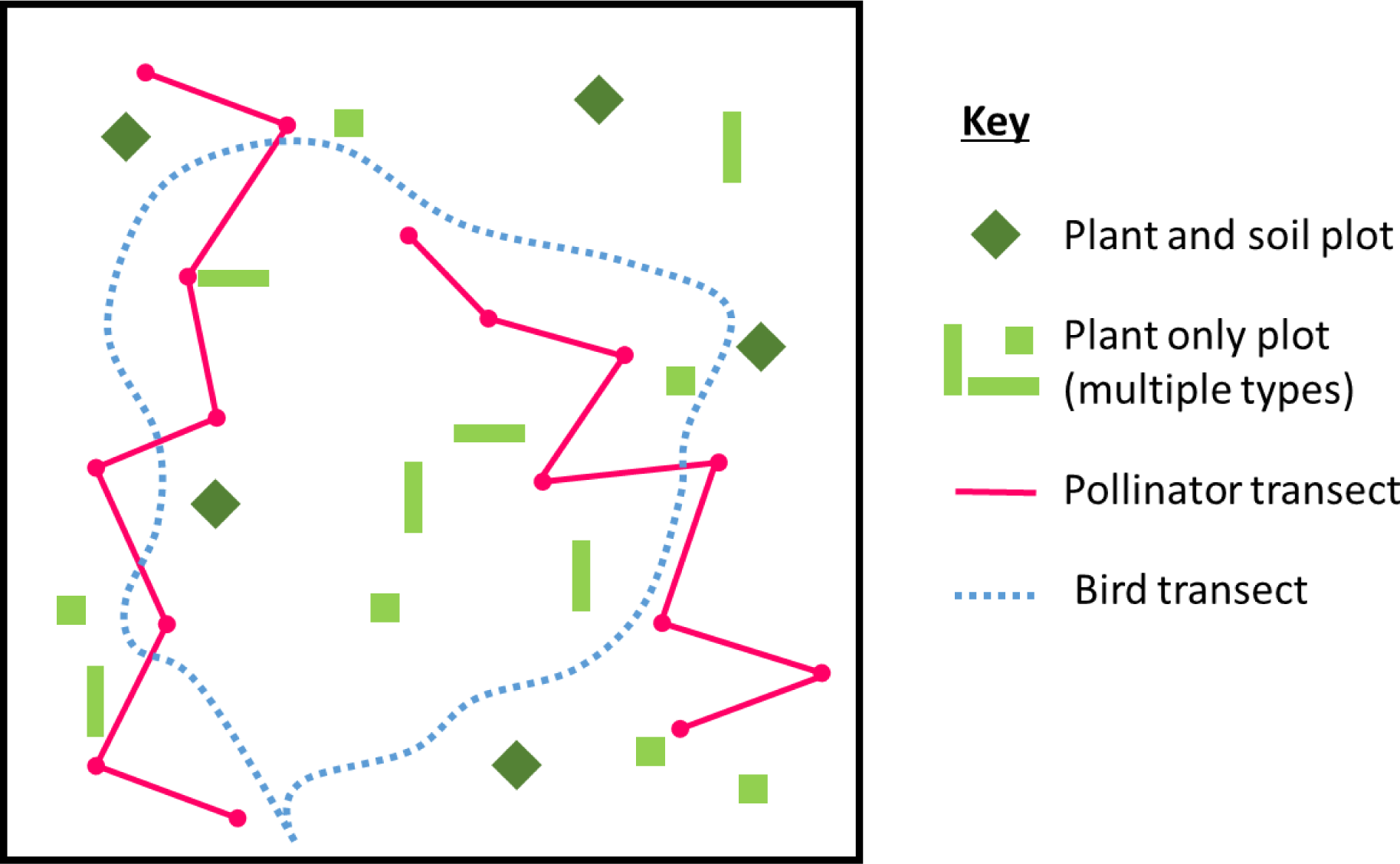
A representative schematic showing the layout of the vegetation plots and pollinator and bird transects within a 1km square. There are 5 randomly located plots with both plant and soil surveyed in each square, while the number and location of plant only plots varies by square as some plot types are habitat specific (e.g. hedgerow plots). Size of plots and transect sections is not to scale.

**Supplementary figure 2.**
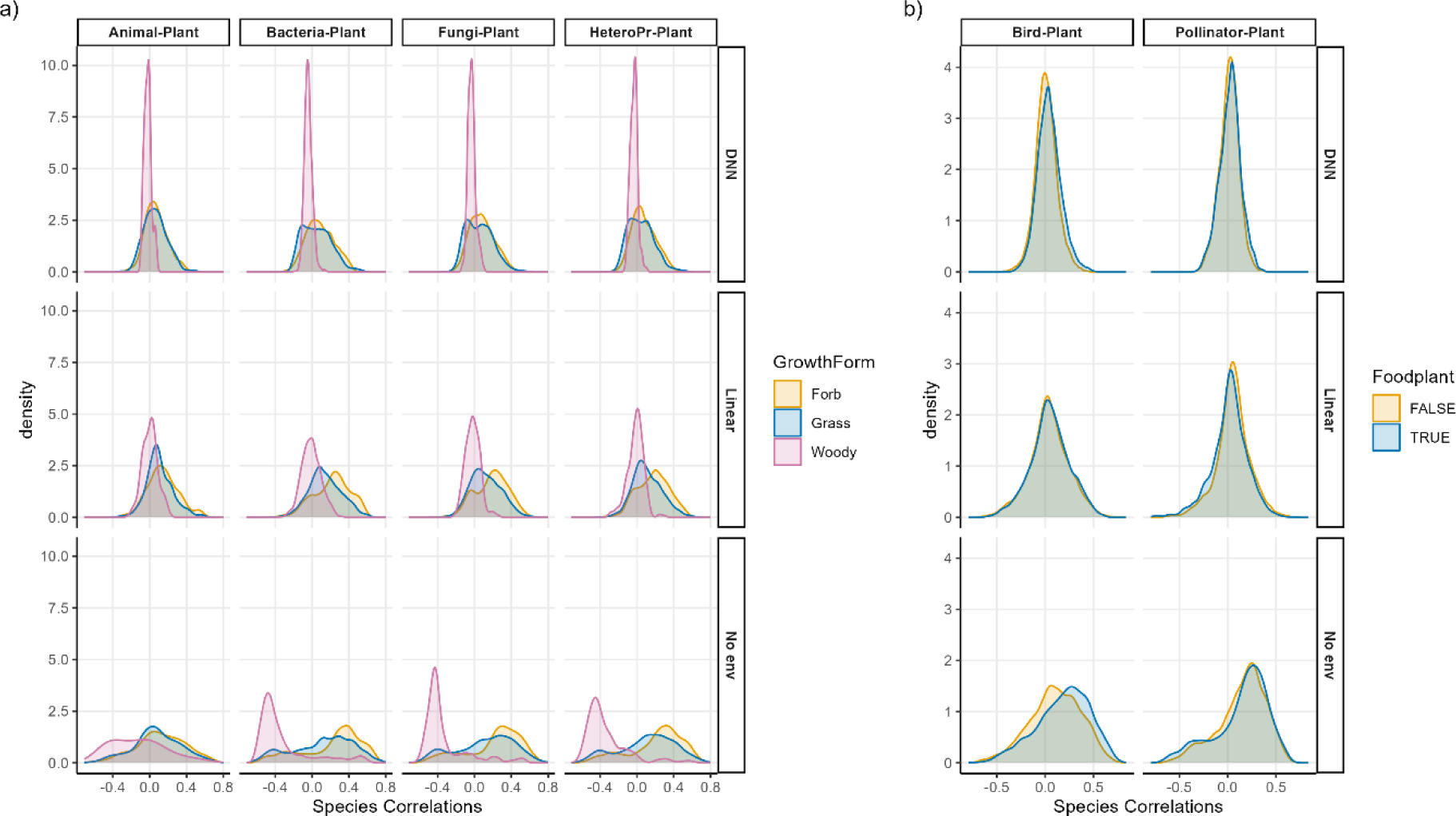
The density distributions of species correlations from the belowground/plot-level (a) and aboveground/square level (b) joint species distribution models that took no account of environment (No env), linear functions of environment only (Linear), or that used deep neural nets to allow for non-linear effects of environment (DNN). Residual correlations are split by plant growth form for the belowground analysis (a) and split by whether the plants are food plants for birds or pollinators for the aboveground analysis (b). Note that despite butterflies, bees and hoverflies being combined into a pollinator category for graphical simplicity the food plants for butterflies are those plants that are important for butterfly larvae while the food plants for bees and hoverflies are those that produce nectar. Also note that there is only woody plant species that appeared in enough plots to be included in the belowground analysis (Vaccinium myrtillus), so those results are unlikely to be representative of all woody species.

**Supplementary figure 3.**
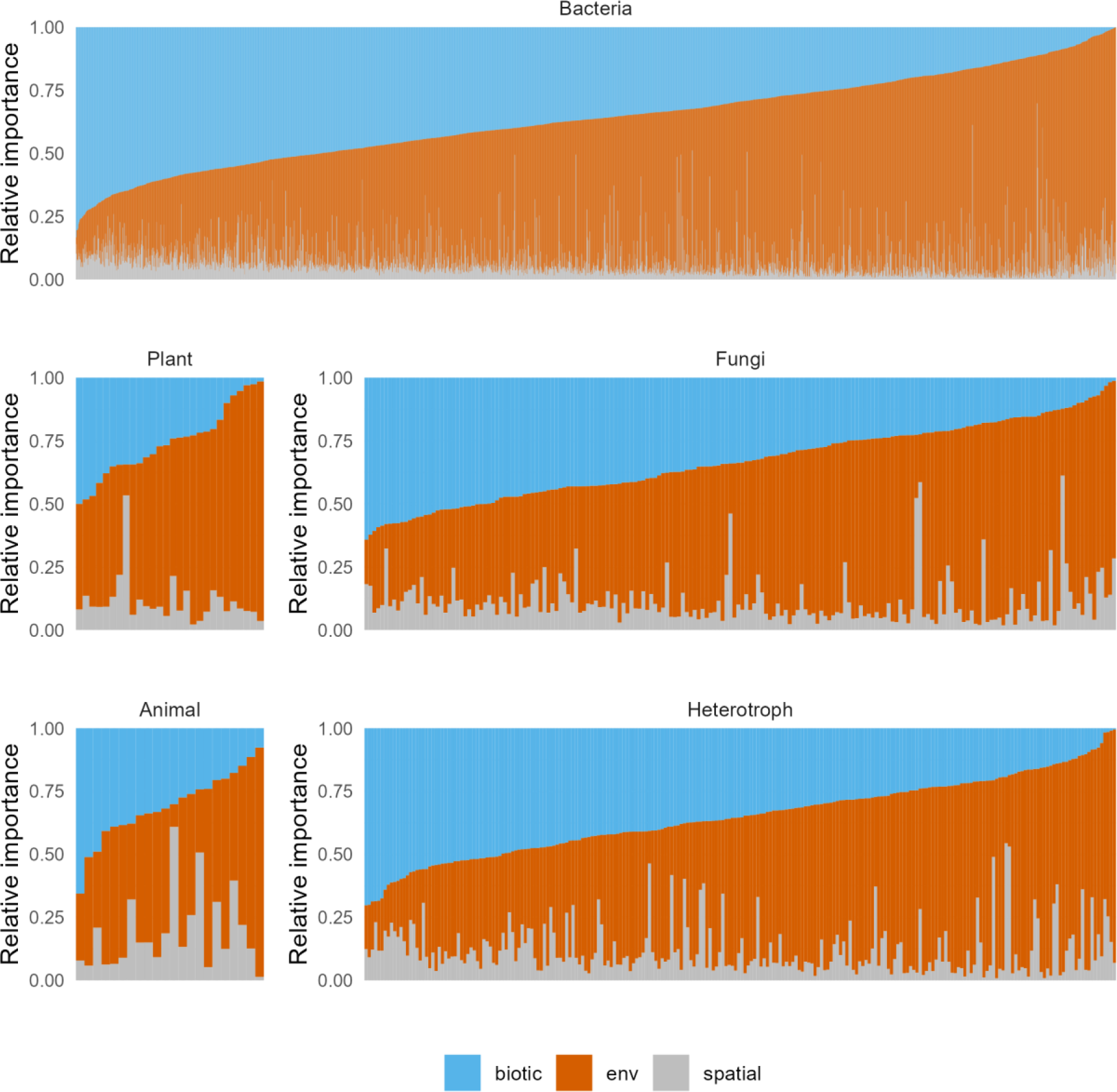
The relative importance (y axis) of the different joint species distribution model components for every bacterial, plant, fungal, animal, and heterotrophic protist taxa (x axis). The importance of the biotic interactions is shown in blue, soil variables (carbon, nitrogen, pH, conductivity, organic matter, water repellency, Olsen-P phosphorus, water content, bulk density) is shown in orange and spatial variation is shown in grey. Taxa are shown in order of decreasing contribution of the biotic component. Results shown are for the model with linear effects of the environment only.

**Supplementary figure 4.**
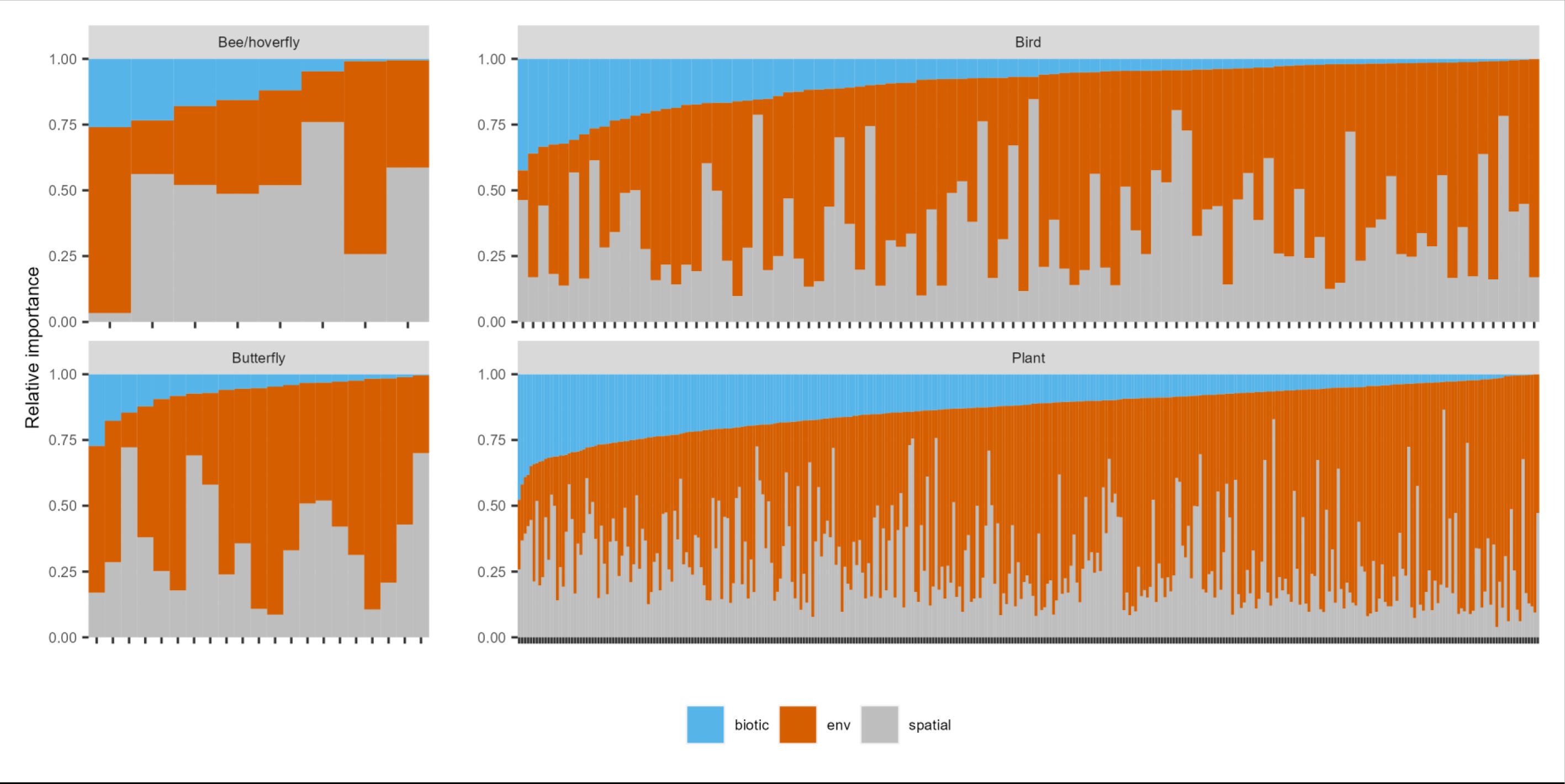
The relative importance (y axis) of the different joint species distribution model components for every bee/hoverfly, bird, butterfly and plant species (x axis). The importance of biotic interactions is shown in blue, the environment (rainfall, temperature and elevation) is shown in orange and spatial variation is shown in grey. Taxa are shown in order of decreasing contribution of the biotic component. Results shown are for the model with linear effects of the environment only.

**Supplementary table 1:** Model parameter estimates from belowground no-environment model.

| Parameter | Mean | Est.Error | l-95% CI | u-95% CI | Rhat | Bulk_ESS | Tail_SS |
| --- | --- | --- | --- | --- | --- | --- | --- |
| **Group-level effects** |  |  |  |  |  |  |  |
| sd(Bacteria) | 477.66 | 34.57 | 413.59 | 549.56 | 1.00 | 5254 | 9557 |
| sd(Fungi) | 50.15 | 5.48 | 39.98 | 61.53 | 1.00 | 3875 | 7184 |
| sd(AM Fungi) | 2.28 | 0.58 | 0.91 | 3.28 | 1.00 | 2847 | 2653 |
| sd(Heterotrophs) | 36.65 | 7.52 | 21.08 | 50.81 | 1.00 | 1976 | 2852 |
| sd(Animals) | 17.83 | 1.79 | 14.41 | 21.50 | 1.00 | 5863 | 8827 |
| sd(Plants) | 2.17 | 0.38 | 1.37 | 2.88 | 1.00 | 3704 | 5497 |
| **Population-level effects** |  |  |  |  |  |  |  |
| Bacteria Intercept | 2307.14 | 44.22 | 2220.48 | 2395.41 | 1.00 | 7659 | 9978 |
| Fungi Intercept | 342.62 | 6.20 | 330.51 | 354.88 | 1.00 | 16093 | 12548 |
| AM Fungi Intercept | 10.06 | 0.39 | 9.29 | 10.81 | 1.00 | 22616 | 13035 |
| Heterotroph Intercept | 277.93 | 5.97 | 266.31 | 289.59 | 1.00 | 22786 | 10836 |
| Animal Intercept | 64.27 | 1.88 | 60.55 | 67.93 | 1.00 | 15502 | 12845 |
| Plant Intercept | 10.54 | 0.29 | 9.98 | 11.10 | 1.00 | 20980 | 12641 |
| **Family specific parameters** |  |  |  |  |  |  |  |
| Sigma Bacteria | 435.93 | 23.67 | 392.12 | 484.28 | 1.00 | 6285 | 9714 |
| Sigma Fungi | 92.01 | 4.82 | 82.68 | 101.66 | 1.00 | 5046 | 8322 |
| Sigma AM Fungi | 6.94 | 0.31 | 6.35 | 7.56 | 1.00 | 6818 | 8536 |
| Sigma Heterotrophs | 101.76 | 5.34 | 91.32 | 112.19 | 1.00 | 3529 | 6672 |
| Sigma Animal | 24.50 | 1.06 | 22.50 | 26.64 | 1.00 | 11884 | 12125 |
| Sigma Plants | 4.59 | 0.21 | 4.20 | 5.03 | 1.00 | 6767 | 10143 |
| **Residual Correlations** |  |  |  |  |  |  |  |
| Bacteria, Fungi | 0.60 | 0.04 | 0.51 | 0.68 | 1.00 | 7008 | 10831 |
| Bacteria, AM Fungi | 0.37 | 0.05 | 0.26 | 0.46 | 1.00 | 13007 | 12551 |
| Fungi, AM Fungi | 0.56 | 0.04 | 0.48 | 0.63 | 1.00 | 19891 | 14261 |
| Bacteria, Heterotrophs | 0.67 | 0.04 | 0.59 | 0.74 | 1.00 | 7798 | 10733 |
| Fungi, Heterotrophs | 0.80 | 0.02 | 0.75 | 0.84 | 1.00 | 8596 | 10582 |
| AM Fungi, Heterotrophs | 0.52 | 0.04 | 0.43 | 0.59 | 1.00 | 21013 | 13808 |
| Bacteria, Animals | 0.24 | 0.06 | 0.12 | 0.35 | 1.00 | 16516 | 13739 |
| Fungi, Animal | 0.24 | 0.06 | 0.12 | 0.35 | 1.00 | 17571 | 13457 |
| AM Fungi, Animal | -0.01 | 0.06 | -0.13 | 0.11 | 1.00 | 17916 | 13082 |
| Heterotroph, Animal | 0.19 | 0.06 | 0.07 | 0.31 | 1.00 | 17929 | 13777 |
| Bacteria, Plant | 0.27 | 0.06 | 0.16 | 0.38 | 1.00 | 18851 | 13946 |
| Fungi, Plant | 0.14 | 0.06 | 0.02 | 0.25 | 1.00 | 19447 | 13581 |
| AM Fungi, Plant | 0.31 | 0.05 | 0.20 | 0.40 | 1.00 | 18497 | 12706 |
| Heterotroph, Plant | 0.17 | 0.06 | 0.06 | 0.28 | 1.00 | 21841 | 13432 |
| Animal, Plant | -0.01 | 0.06 | -0.12 | 0.11 | 1.00 | 18000 | 13017 |

**Supplementary table 2:** Model parameter estimates from belowground linear environment model.

| Parameter | Mean | Est.Error | l-95% CI | u-95% CI | Rhat | Bulk_ESS | Tail_SS |
| --- | --- | --- | --- | --- | --- | --- | --- |
| **Group-level effects** |  |  |  |  |  |  |  |
| sd(Bacteria) | 485.09 | 35.34 | 420.42 | 558.35 | 1.00 | 4365 | 8353 |
| sd(Fungi) | 51.58 | 5.69 | 41.09 | 63.34 | 1.00 | 2639 | 5804 |
| sd(AM Fungi) | 2.39 | 0.52 | 1.23 | 3.29 | 1.00 | 2502 | 2074 |
| sd(Heterotrophs) | 35.91 | 8.7 | 16.53 | 51.76 | 1.00 | 1323 | 1465 |
| sd(Animals) | 13.93 | 1.75 | 10.53 | 17.44 | 1.00 | 4820 | 8324 |
| sd(Plants) | 2.15 | 0.36 | 1.41 | 2.83 | 1.00 | 4075 | 5802 |
| **Population-level effects** |  |  |  |  |  |  |  |
| Bacteria Intercept | 2308.10 | 45.41 | 2218.16 | 2396.38 | 1.00 | 5004 | 7864 |
| Fungi Intercept | 347.58 | 8.91 | 330.37 | 364.81 | 1.00 | 15531 | 11916 |
| AM Fungi Intercept | 10.07 | 2.35 | 5.40 | 14.68 | 1.00 | 18448 | 12795 |
| Heterotroph Intercept | 285.12 | 8.69 | 268.24 | 302.26 | 1.00 | 21240 | 11439 |
| Animal Intercept | 51.25 | 5.52 | 40.56 | 62.15 | 1.00 | 23026 | 11874 |
| Plant Intercept | 5.86 | 1.97 | 1.98 | 9.68 | 1.00 | 15542 | 11882 |
| Bacteria pH | 0.24 | 1.00 | -1.72 | 2.20 | 1.00 | 31809 | 11227 |
| Bacteria Carbon | -0.12 | 1.00 | -2.08 | 1.83 | 1.00 | 30469 | 11496 |
| Bacteria Water | -0.01 | 0.98 | -1.93 | 1.88 | 1.00 | 28298 | 11831 |
| Bacteria Total P | -0.05 | 0.99 | -1.98 | 1.88 | 1.00 | 32645 | 11249 |
| Bacteria SWR | -0.25 | 0.99 | -2.18 | 1.68 | 1.00 | 29401 | 11595 |
| Fungi pH | 0.49 | 0.98 | -1.43 | 2.41 | 1.00 | 28923 | 11225 |
| Fungi Carbon | -1.28 | 0.98 | -3.23 | 0.65 | 1.00 | 27564 | 11324 |
| Fungi Water | -0.25 | 1.01 | -2.22 | 1.73 | 1.00 | 31189 | 11011 |
| Fungi Total P | -0.27 | 0.99 | -2.23 | 1.65 | 1.00 | 28801 | 11403 |
| Fungi SWR | -0.86 | 0.84 | -2.53 | 0.81 | 1.00 | 22227 | 11631 |
| AM Fungi pH | 0.54 | 0.40 | -0.24 | 1.31 | 1.00 | 16135 | 12384 |
| AM Fungi Carbon | -2.02 | 0.44 | -2.86 | -1.16 | 1.00 | 13846 | 12595 |
| AM Fungi Water | -0.05 | 0.91 | -1.83 | 1.73 | 1.00 | 26581 | 11602 |
| AM Fungi Total P | 1.40 | 0.55 | 0.32 | 2.47 | 1.00 | 17351 | 12966 |
| AM Fungi SWR | 0.33 | 0.14 | 0.05 | 0.60 | 1.00 | 17349 | 12711 |
| Heterotrophs pH | 1.42 | 0.99 | -0.52 | 3.37 | 1.00 | 27496 | 11069 |
| Heterotrophs Carbon | -1.02 | 0.98 | -2.93 | 0.90 | 1.00 | 29679 | 11271 |
| Heterotrophs Water | -0.07 | 0.99 | -2.01 | 1.87 | 1.00 | 27640 | 10821 |
| Heterotrophs Total P | 0.54 | 1.00 | -1.43 | 2.50 | 1.00 | 29504 | 11148 |
| Heterotrophs SWR | -2.21 | 0.87 | -3.90 | -0.52 | 1.00 | 23872 | 11840 |
| Animal pH | -2.09 | 0.89 | -3.85 | -0.31 | 1.00 | 21494 | 12070 |
| Animal Carbon | 3.87 | 0.90 | 2.10 | 5.61 | 1.00 | 23362 | 12676 |
| Animal Water | 0.78 | 1.01 | -1.21 | 2.75 | 1.00 | 29958 | 11553 |
| Animal Total P | 0.08 | 0.94 | -1.78 | 1.91 | 1.00 | 26089 | 11870 |
| Animal SWR | 2.72 | 0.5 | 1.74 | 3.69 | 1.00 | 19709 | 11363 |
| Plant pH | 1.12 | 0.34 | 0.48 | 1.79 | 1.00 | 15392 | 11842 |
| Plant Carbon | -0.71 | 0.37 | -1.42 | 0.01 | 1.00 | 14657 | 11767 |
| Plant Water | 1.02 | 0.86 | -0.69 | 2.71 | 1.00 | 25028 | 12902 |
| Plant Total P | -0.56 | 0.46 | -1.48 | 0.36 | 1.00 | 18584 | 12316 |
| Plant SWR | 0.15 | 0.12 | -0.08 | 0.38 | 1.00 | 17736 | 12660 |
| **Family specific parameters** |  |  |  |  |  |  |  |
| Sigma Bacteria | 429.28 | 23.30 | 386.31 | 476.99 | 1.00 | 4138 | 7342 |
| Sigma Fungi | 89.27 | 4.95 | 79.82 | 99.15 | 1.00 | 3043 | 6223 |
| Sigma AM Fungi | 6.14 | 0.30 | 5.58 | 6.76 | 1.00 | 5698 | 7606 |
| Sigma Heterotrophs | 98.32 | 5.67 | 87.5 | 109.68 | 1.00 | 1933 | 3557 |
| Sigma Animal | 24.71 | 1.10 | 22.68 | 26.93 | 1.00 | 8999 | 11458 |
| Sigma Plants | 4.36 | 0.20 | 3.99 | 4.77 | 1.00 | 6737 | 9422 |
| **Residual Correlations** |  |  |  |  |  |  |  |
| Bacteria, Fungi | 0.58 | 0.05 | 0.48 | 0.67 | 1.00 | 4469 | 8046 |
| Bacteria, AM Fungi | 0.27 | 0.06 | 0.15 | 0.39 | 1.00 | 9362 | 11687 |
| Fungi, AM Fungi | 0.46 | 0.05 | 0.36 | 0.56 | 1.00 | 12405 | 11997 |
| Bacteria, Heterotrophs | 0.65 | 0.04 | 0.56 | 0.72 | 1.00 | 4542 | 8129 |
| Fungi, Heterotrophs | 0.78 | 0.03 | 0.73 | 0.83 | 1.00 | 4276 | 7634 |
| AM Fungi, Heterotrophs | 0.41 | 0.06 | 0.29 | 0.51 | 1.00 | 11636 | 12142 |
| Bacteria, Animals | 0.41 | 0.05 | 0.30 | 0.51 | 1.00 | 10812 | 12481 |
| Fungi, Animal | 0.43 | 0.05 | 0.32 | 0.53 | 1.00 | 12233 | 11743 |
| AM Fungi, Animal | 0.12 | 0.06 | 0.00 | 0.23 | 1.00 | 16028 | 13084 |
| Heterotroph, Animal | 0.40 | 0.06 | 0.28 | 0.5 | 1.00 | 12937 | 12049 |
| Bacteria, Plant | 0.12 | 0.07 | -0.02 | 0.25 | 1.00 | 12923 | 13151 |
| Fungi, Plant | -0.02 | 0.07 | -0.15 | 0.12 | 1.00 | 13761 | 12593 |
| AM Fungi, Plant | 0.21 | 0.06 | 0.09 | 0.32 | 1.00 | 13550 | 12713 |
| Heterotroph, Plant | 0.01 | 0.07 | -0.13 | 0.15 | 1.00 | 12678 | 12114 |
| Animal, Plant | 0.02 | 0.06 | -0.09 | 0.14 | 1.00 | 17748 | 12603 |

**Supplementary table 3:** Model parameter estimates from belowground sigmoidal pH model, where richness ∼ delta + alpha/(1 + exp(-beta*(pH – gamma))

| Parameter | Mean | Est.Error | l-95% CI | u-95% CI | Rhat | Bulk_ESS | Tail_SS |
| --- | --- | --- | --- | --- | --- | --- | --- |
| **Group-level effects** |  |  |  |  |  |  |  |
| sd(Bacteria) | 483.85 | 32.61 | 424.03 | 551.00 | 1.00 | 4819 | 8526 |
| sd(Fungi) | 48.89 | 5.00 | 39.42 | 59.01 | 1.00 | 4652 | 8230 |
| sd(AM Fungi) | 2.35 | 0.47 | 1.34 | 3.20 | 1.00 | 3846 | 4596 |
| sd(Heterotrophs) | 36.15 | 5.80 | 24.51 | 47.32 | 1.00 | 3450 | 5180 |
| sd(Animals) | 13.91 | 1.90 | 10.12 | 17.66 | 1.00 | 4880 | 7739 |
| sd(Plants) | 1.90 | 0.38 | 1.09 | 2.59 | 1.00 | 3367 | 4043 |
| **Population-level effects** |  |  |  |  |  |  |  |
| Bacteria delta | -815.07 | 504.36 | -1810.22 | 180.76 | 1.00 | 12744 | 11036 |
| Bacteria alpha | 3729.08 | 500.27 | 2740.88 | 4711.39 | 1.00 | 13426 | 11601 |
| Bacteria beta | 0.85 | 0.14 | 0.60 | 1.13 | 1.00 | 13886 | 12847 |
| Bacteria gamma | 2.21 | 0.31 | 1.62 | 2.86 | 1.00 | 11008 | 10699 |
| Fungi delta | 187.06 | 25.97 | 131.31 | 232.50 | 1.00 | 9484 | 9775 |
| Fungi alpha | 220.03 | 29.32 | 167.45 | 282.80 | 1.00 | 9839 | 11044 |
| Fungi beta | 1.84 | 0.35 | 1.25 | 2.60 | 1.00 | 13547 | 11455 |
| Fungi gamma | 3.65 | 0.18 | 3.28 | 3.98 | 1.00 | 11472 | 10999 |
| AM Fungi delta | 0.77 | 1.69 | -3.20 | 3.12 | 1.00 | 7903 | 5083 |
| AM Fungi alpha | 12.73 | 1.76 | 10.17 | 16.88 | 1.00 | 8319 | 5456 |
| AM Fungi beta | 3.37 | 0.59 | 2.24 | 4.56 | 1.00 | 9362 | 6480 |
| AM Fungi gamma | 3.70 | 0.14 | 3.37 | 3.92 | 1.00 | 8174 | 5392 |
| Heterotrophs delta | 40.24 | 45.66 | -55.61 | 119.52 | 1.00 | 7675 | 10751 |
| Heterotrophs alpha | 342.78 | 50.95 | 249.80 | 447.29 | 1.00 | 8057 | 11253 |
| Heterotrophs beta | 1.38 | 0.28 | 0.95 | 2.06 | 1.00 | 9392 | 11202 |
| Heterotrophs gamma | 3.59 | 0.23 | 3.10 | 4.00 | 1.00 | 7968 | 10348 |
| Animal delta | 44.81 | 7.24 | 29.88 | 57.68 | 1.00 | 13041 | 12309 |
| Animal alpha | 46.50 | 10.16 | 26.82 | 66.37 | 1.00 | 15245 | 9742 |
| Animal beta | -0.62 | 0.33 | -1.44 | -0.18 | 1.00 | 11085 | 8079 |
| Animal gamma | 3.80 | 0.92 | 2.04 | 5.67 | 1.00 | 19688 | 12257 |
| Plant delta | -3.68 | 6.46 | -19.27 | 4.90 | 1.00 | 8989 | 9031 |
| Plant alpha | 15.8 | 6.49 | 7.10 | 31.42 | 1.00 | 8944 | 9041 |
| Plant beta | 2.08 | 0.53 | 1.22 | 3.29 | 1.00 | 13516 | 12529 |
| Plant gamma | 2.77 | 0.39 | 1.98 | 3.44 | 1.00 | 9098 | 10654 |
| **Family specific parameters** |  |  |  |  |  |  |  |
| Sigma Bacteria | 324.95 | 15.71 | 295.73 | 356.94 | 1.00 | 9070 | 12594 |
| Sigma Fungi | 74.92 | 3.65 | 68.09 | 82.43 | 1.00 | 6450 | 10399 |
| Sigma AM Fungi | 5.63 | 0.25 | 5.16 | 6.14 | 1.00 | 7722 | 8874 |
| Sigma Heterotrophs | 76.35 | 3.86 | 69.2 | 84.32 | 1.00 | 5292 | 7938 |
| Sigma Animal | 25.66 | 1.16 | 23.48 | 28.05 | 1.00 | 8109 | 11683 |
| Sigma Plants | 4.29 | 0.19 | 3.94 | 4.68 | 1.00 | 6650 | 7518 |
| **Residual Correlations** |  |  |  |  |  |  |  |
| Bacteria, Fungi | 0.38 | 0.06 | 0.26 | 0.48 | 1.00 | 8392 | 11520 |
| Bacteria, AM Fungi | 0.06 | 0.06 | -0.06 | 0.17 | 1.00 | 15094 | 13628 |
| Fungi, AM Fungi | 0.36 | 0.05 | 0.25 | 0.45 | 1.00 | 16897 | 13100 |
| Bacteria, Heterotrophs | 0.42 | 0.05 | 0.31 | 0.52 | 1.00 | 8724 | 12094 |
| Fungi, Heterotrophs | 0.68 | 0.03 | 0.61 | 0.74 | 1.00 | 9808 | 12968 |
| AM Fungi, Heterotrophs | 0.26 | 0.05 | 0.16 | 0.37 | 1.00 | 18659 | 12906 |
| Bacteria, Animals | 0.35 | 0.05 | 0.24 | 0.45 | 1.00 | 12776 | 13129 |
| Fungi, Animal | 0.36 | 0.05 | 0.25 | 0.46 | 1.00 | 13437 | 13853 |
| AM Fungi, Animal | 0.03 | 0.06 | -0.08 | 0.14 | 1.00 | 20372 | 13474 |
| Heterotroph, Animal | 0.34 | 0.05 | 0.23 | 0.44 | 1.00 | 14176 | 12433 |
| Bacteria, Plant | 0.06 | 0.06 | -0.05 | 0.17 | 1.00 | 18625 | 13447 |
| Fungi, Plant | -0.10 | 0.06 | -0.21 | 0.01 | 1.00 | 18670 | 13472 |
| AM Fungi, Plant | 0.12 | 0.05 | 0.02 | 0.23 | 1.00 | 21454 | 13134 |
| Heterotroph, Plant | -0.10 | 0.06 | -0.21 | 0.02 | 1.00 | 20854 | 13135 |
| Animal, Plant | 0.00 | 0.06 | -0.11 | 0.11 | 1.00 | 21656 | 13670 |

**Supplementary table 4:** Model parameter estimates for the square-level model with no environmental variables.

| Parameter | Mean | Est.Error | l-95% CI | u-95% CI | Rhat | Bulk_ESS | Tail_SS |
| --- | --- | --- | --- | --- | --- | --- | --- |
| **Group-level effects** |  |  |  |  |  |  |  |
| sd(Bird) | 0.19 | 0.06 | 0.10 | 0.33 | 1.00 | 4778 | 7544 |
| sd(Butterfly) | 0.18 | 0.06 | 0.07 | 0.32 | 1.00 | 3436 | 3102 |
| sd(Bee) | 0.13 | 0.03 | 0.08 | 0.19 | 1.00 | 6108 | 9760 |
| sd(Plant) | 0.04 | 0.03 | 0.00 | 0.12 | 1.00 | 5522 | 6382 |
| **Population-level effects** |  |  |  |  |  |  |  |
| Bird Intercept | 2.87 | 0.06 | 2.75 | 2.99 | 1.00 | 5834 | 7708 |
| Butterfly Intercept | 1.31 | 0.06 | 1.20 | 1.43 | 1.00 | 6805 | 9244 |
| Bee Intercept | 0.37 | 0.04 | 0.29 | 0.44 | 1.00 | 4657 | 7232 |
| Plant Intercept | 2.95 | 0.03 | 2.90 | 3.01 | 1.00 | 13772 | 10780 |
| **Family specific parameters** |  |  |  |  |  |  |  |
| Sigma Bird | 0.60 | 0.03 | 0.55 | 0.65 | 1.00 | 11991 | 11968 |
| Sigma Butterfly | 0.55 | 0.02 | 0.51 | 0.60 | 1.00 | 11464 | 11140 |
| Sigma Bee | 0.31 | 0.01 | 0.28 | 0.33 | 1.00 | 15334 | 12013 |
| Sigma Plant | 0.44 | 0.02 | 0.40 | 0.48 | 1.00 | 17887 | 12348 |
| **Residual Correlations** |  |  |  |  |  |  |  |
| Bird, Butterfly | 0.44 | 0.05 | 0.35 | 0.54 | 1.00 | 12775 | 11460 |
| Bird, Bee | 0.40 | 0.05 | 0.30 | 0.50 | 1.00 | 12517 | 12002 |
| Butterfly, Bee | 0.40 | 0.05 | 0.30 | 0.50 | 1.00 | 14866 | 12384 |
| Bird, Plant | 0.41 | 0.05 | 0.31 | 0.50 | 1.00 | 16577 | 12828 |
| Butterfly, Plant | 0.27 | 0.06 | 0.16 | 0.38 | 1.00 | 16249 | 12317 |
| Bee, Plant | 0.24 | 0.06 | 0.13 | 0.35 | 1.00 | 16463 | 12440 |

**Supplementary table 5:** Model parameter estimates for the square-level model with linear environmental effects.

| Parameter | Mean | Est.Error | l-95% CI | u-95% CI | Rhat | Bulk_ESS | Tail_SS |
| --- | --- | --- | --- | --- | --- | --- | --- |
| **Group-level effects** |  |  |  |  |  |  |  |
| sd(Bird) | 0.07 | 0.04 | 0.00 | 0.16 | 1.00 | 4906 | 7476 |
| sd(Butterfly) | 0.08 | 0.04 | 0.00 | 0.17 | 1.00 | 4597 | 7665 |
| sd(Bee) | 0.13 | 0.03 | 0.08 | 0.19 | 1.00 | 7743 | 10212 |
| sd(Plant) | 0.07 | 0.04 | 0.00 | 0.16 | 1.00 | 4496 | 6900 |
| **Population-level effects** |  |  |  |  |  |  |  |
| Bird Intercept | 2.87 | 0.03 | 2.80 | 2.94 | 1.00 | 18563 | 11184 |
| Butterfly Intercept | 1.31 | 0.04 | 1.24 | 1.38 | 1.00 | 17306 | 11421 |
| Bee Intercept | 0.36 | 0.04 | 0.28 | 0.42 | 1.00 | 9322 | 10928 |
| Plant Intercept | 2.95 | 0.03 | 2.89 | 3.02 | 1.00 | 18104 | 11705 |
| Bird Rainfall | -0.16 | 0.04 | -0.23 | -0.08 | 1.00 | 17187 | 13273 |
| Bird Temperature | 0.28 | 0.04 | 0.20 | 0.34 | 1.00 | 18637 | 12865 |
| Butterfly Rainfall | -0.21 | 0.04 | -0.28 | -0.13 | 1.00 | 17696 | 11189 |
| Butterfly Temperature | 0.17 | 0.04 | 0.10 | 0.24 | 1.00 | 17741 | 12927 |
| Bee Rainfall | -0.04 | 0.03 | -0.09 | 0.01 | 1.00 | 16468 | 12460 |
| Bee Temperature | 0.14 | 0.02 | 0.09 | 0.19 | 1.00 | 16750 | 12887 |
| Plant Rainfall | -0.02 | 0.03 | -0.09 | 0.04 | 1.00 | 19208 | 13308 |
| Plant Temperature | 0.14 | 0.03 | 0.08 | 0.21 | 1.00 | 13754 | 11802 |
| **Family specific parameters** |  |  |  |  |  |  |  |
| Sigma Bird | 0.46 | 0.02 | 0.42 | 0.50 | 1.00 | 28206 | 12267 |
| Sigma Butterfly | 0.45 | 0.02 | 0.41 | 0.49 | 1.00 | 24489 | 12021 |
| Sigma Bee | 0.26 | 0.01 | 0.24 | 0.28 | 1.00 | 24091 | 12640 |
| Sigma Plant | 0.41 | 0.02 | 0.37 | 0.44 | 1.00 | 22690 | 12417 |
| **Residual Correlations** |  |  |  |  |  |  |  |
| Bird, Butterfly | 0.11 | 0.06 | 0.00 | 0.23 | 1.00 | 28912 | 11151 |
| Bird, Bee | 0.10 | 0.06 | -0.02 | 0.22 | 1.00 | 29749 | 11640 |
| Butterfly, Bee | 0.15 | 0.06 | 0.03 | 0.26 | 1.00 | 27279 | 11895 |
| Bird, Plant | 0.24 | 0.06 | 0.13 | 0.34 | 1.00 | 27174 | 11776 |
| Butterfly, Plant | 0.07 | 0.06 | -0.04 | 0.19 | 1.00 | 30723 | 11240 |
| Bee, Plant | 0.04 | 0.06 | -0.08 | 0.16 | 1.00 | 31735 | 11764 |

**Supplementary table 6:**
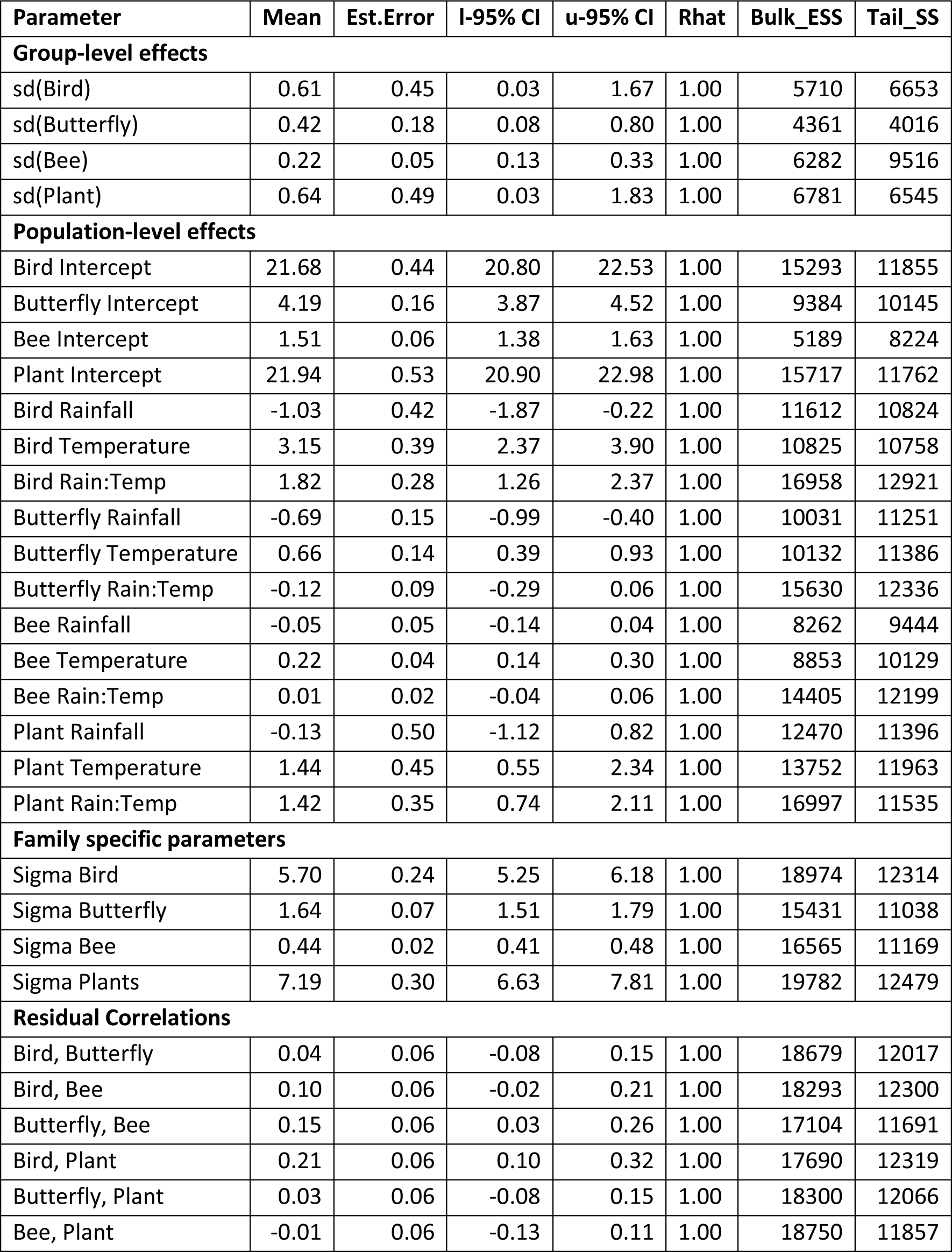
Model parameter estimates from aboveground temperature*rainfall model to exponentiated diversity values.

## Notes

### Competing Interest Statement

The authors have declared no competing interest.

### Summary of Updates

Minor revisions to the text, in particular providing more justification for our environmental gradient choices in the methods and adding more context to the discussion.

